# A New Approach to Testing Mediation of the Microbiome using the LDM

**DOI:** 10.1101/2021.11.12.468449

**Authors:** Ye Yue, Yi-Juan Hu

## Abstract

**Background:** Understanding whether and which microbes played a mediating role between an exposure and a disease outcome are essential for researchers to develop clinical interventions to treat the disease by modulating the microbes. Existing methods for mediation analysis of the microbiome are often limited to a global test of community-level mediation or selection of mediating microbes without control of the false discovery rate (FDR). Further, while the null hypothesis of no mediation at each microbe is a composite null that consists of three types of null (no exposure-microbe association, no microbe-outcome association given the exposure, or neither), most existing methods for the global test such as MedTest and MODIMA treat the microbes as if they are all under the same type of null.

**Methods:** We propose a new approach based on *inverse regression* that regresses the (possibly transformed) relative abundance of each taxon on the exposure and the exposure-adjusted outcome to assess the exposure-taxon and taxon-outcome associations simultaneously. Then the association *p*-values are used to test mediation at both the community and individual taxon levels. This approach fits nicely into our Linear Decomposition Model (LDM) framework, so our new method is implemented in the LDM and enjoys all the features of the LDM, i.e., allowing an arbitrary number of taxa to be tested, supporting continuous, discrete, or multivariate exposures and outcomes as well as adjustment of confounding covariates, accommodating clustered data, and offering analysis at the relative abundance or presence-absence scale. We refer to this new method as LDM-med.

**Results:** Using extensive simulations, we showed that LDM-med always controlled the type I error of the global test and had compelling power over existing methods; LDM-med always preserved the FDR of testing individual taxa and had much better sensitivity than alternative approaches. In contrast, MedTest and MODIMA had severely inflated type I error when different taxa were under different types of null. The flexibility of LDM-med for a variety of mediation analyses is illustrated by the application to a murine microbiome dataset.

**Availability and Implementation:** Our new method has been added to our R package LDM, which is available on GitHub at https://github.com/yijuanhu/LDM.

## Background

While most microbiome studies conducted so far have focused on bivariate associations between the microbiome and their covariates of interest [1, 2], increasing studies have emerged recently to elucidate the biological mechanisms underlying the complex interplay between environmental exposures, the microbiome, and clinical outcomes. In many cases, it is of interest to understand whether the microbiome play a mediating role between an exposure and an outcome [3–5], as depicted in Figure 1(a). For example, does diet have some effect on inflammatory bowel diseases that is mediated through the perturbation of the gut microbiome [4]? How does the change in the gut microbiome due to antibiotic exposure cause the change in mouse body weight [5]? It is of particular importance to single out the specific microbes that are responsible for the overall mediation effect, which is essential for researchers to develop clinical interventions to modify the outcome by modulating the mediating microbes [6].

**Figure 1:**
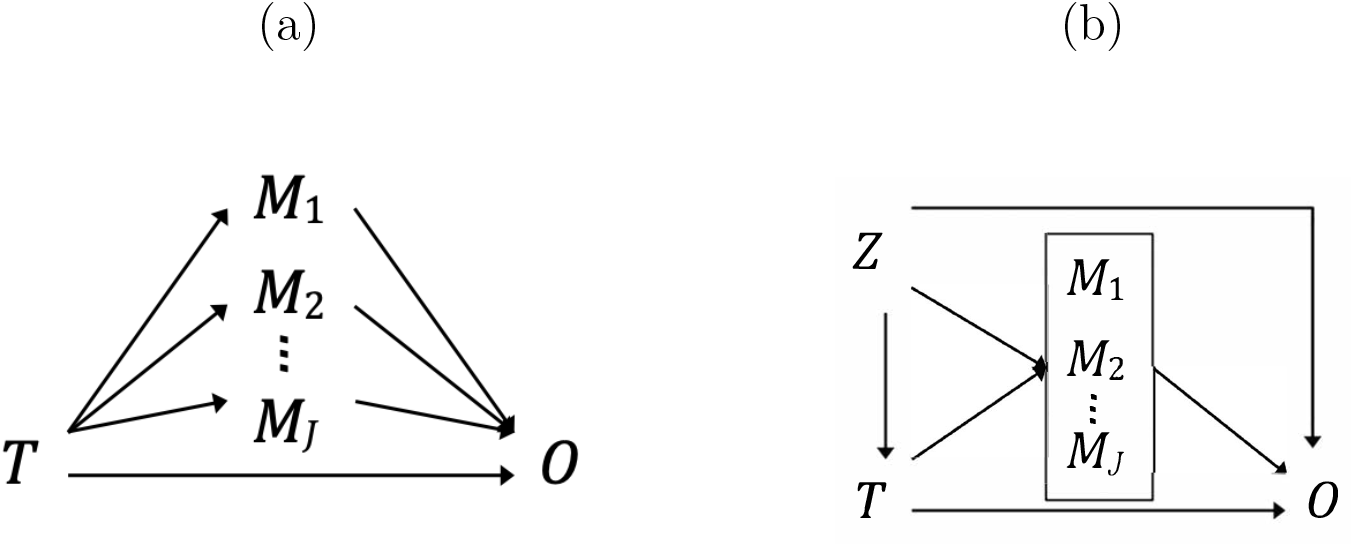
(a) Some effect of the exposure on the outcome is mediated through multiple microbes. (b) *T* denotes the exposure, (*M*_1_,…, *M*_*j*_) the microbes, *O* the outcome, and *Z* the confounding covariates.

Let *T* denote the exposure (treatment), *M* = (*M*_1_,…, *M_J_*) the *J* mediators, *O* the outcome, and *Z* the confounding covariates; using this notation, the mediation relationships are shown in Figure 1(b). To claim a mediation effect of a microbe, both the exposure-microbe and microbe-outcome associations (given the exposure) are required to be significant. Thus, the null hypothesis of no mediation at microbe *j* is a composite null that consists of no microbe-outcome association, no exposure-microbe association, or neither:

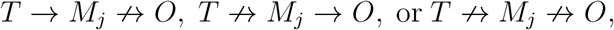

which are referred to as the type-I, type-II, and type-III null hypotheses, respectively. It is highly likely that different microbes are under different types of null. For example, antibiotic use may perturb a large number of microbes but most of them do no modify mouse body weight, whereas some microbes remain intact from antibiotic use but do interact with the body weight; of course, there are microbes that are not associated with either factor. In this example, we have all three types of null.

Read count data from 16S amplicon or metagenomic sequencing are typically summarized in a taxa count table; here we use “taxon” generically to refer to any feature such as operational taxonomic unites, amplicon sequence variants, or any other taxonomic or functional grouping of bacterial sequences. The taxa count data have unique and complex features. They are high-dimensional with typically many more taxa than samples, and it is desirable to test all taxa simultaneously for mediation effects with multiple testing correction that controls the false discovery rate (FDR). The data are also sparse (having 50-90% zero counts), compositional (measuring relative abundances that sum to one), and highly overdispersed. In addition, microbiome studies may have complex exposures or outcomes that can not only be continuous but also discrete, as well as multivariate (comprising multiple components such as categorical variables with more than two levels). These studies also often have small sample sizes (e.g., 50–100) and complex designs (e.g., clustered data [7], matched sets [8], longitudinal sampling). The capability to handle all these features is essential for any statistical method to be practically useful.

Because it is challenging to conduct mediation analysis in the high-dimensional setting, some existing methods are limited to testing the overall mediation effect at the community level, while others attemp to identify mediating taxa but have no control of the FDR. Specifically, MedTest [9] and MODIMA [10] base their tests on distance matrices that summarize the high-dimensional data into between-sample dissimilarity measurements, and thus produce a global *p*-value only. Also, they treat the taxa as if they are all under the same type of null in their permutation procedures that provide the null distributions of the test statistics. Although they can accommodate binary outcomes without any modification, MedTest does not allow multivariate exposures or outcomes, whereas MODIMA does not allow adjustment of confounding covariates. Finally, MedTest, by using the top ten principal components of the distance matrix as ten mediators, may not capture mediation effects in rare taxa (even when the Jaccard distance is used in the omnibus test); MODIMA highly depends on the choice of the distance metric and currently does not provide an omnibus test. CMM [11] (and CMMB, the extension for binary outcomes [12]) and SparseMCMM [5] consider the high-dimensional mediators individually and conduct estimation and testing of mediation effects at both the community level and the taxon level. However, SparseMCMM selects mediating taxa using regularization techniques and CMM using confidence interval estimates, so neither has formal control of the FDR. Also, they are limited to analyzing simple continuous outcomes. Because they add pseudocounts to the (massive) zero counts when performing some log-ratio transformation of the read count data as a way to handle compositionality, they are prone to false positive findings as demonstrated in the context of testing taxon differential abundance[13]. Zhang’s method [14] shares all the features in CMM and SparseMCMM, except that Zheng’s method tests only one taxon in one run, which is the first isometric log-ratio (ilr) transformed variable (because other ilr variables are not interpretable), and is impractical for use with hundreds and thousands of taxa.

In this article, we focus on testing, rather than estimation, of marginal mediation effects at individual taxa with a goal of controlling the FDR. This strategy is very common in the initial *scan* of high-dimensional features in omic studies [15–17]; “fine mapping” of mechanistic mediators and formal estimation of their mediation effects can be performed more easily after the dimension is greatly reduced. We find that, the testing objective can be achieved by utilizing inverse regression and the Linear Decomposition Model (LDM) framework [7, 8, 18, 19] we developed originally for testing microbiome associations [7], which together enable us to handle all the aforementioned data complexities in a natural way. In the methods section, we first describe our method for testing individual taxa for mediation and then a method that aggregates the taxon-level information to test the overall mediation in a community. In the results section, we present extensive simulation results in which we numerically compared our method, which we call LDM-med, to alternative methods, and we apply the new method to data from a murine study. We conclude with a discussion section.

## Methods

### Motivation

Our starting point is the following classical model for multiple mediators [20]. For a continuous outcome and *J* continuous (potential) mediators with no exposure-mediator and mediator-mediator interactions, the model specifies a linear model for each mediator and a linear model for the outcome that includes all mediators:

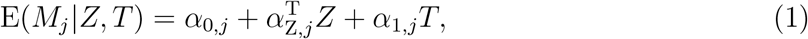

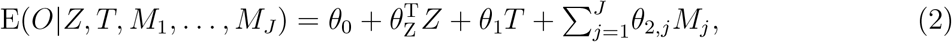

where the notation was introduced in Figure 1(b). It can be derived that the overall (total) mediation effect through (*M*_1_,…, *M_J_*) takes the form 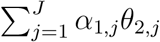 [20]; note that *α*_1,*j*_ characterizes the effect of *T* on *M*_*j*_ given *Z* and *θ*_2,*j*_ the effect of *M*_*j*_ on *O* given *Z* and *T* and all other *M*_*j*_s. When the mediators are independent of one another conditional on *Z* and *T*, each product *α*_1,*j*_*θ*_2,*j*_ can be interpreted as the mediation effect through a single mediator *M*_*j*_. Even if the mediators are not conditionally independent, a non-zero value of *α*_1,*j*_*θ*_2,*j*_ indicates a non-zero contribution of *M*_*j*_ to the overall mediation effect. Thus, our objective can be achieved by testing whether *α*_1,*j*_*θ*_2,*j*_ = 0 at each potential mediator. However, the *forward* outcome model (2), although describing the mediation process in a natural order and enabling intuitive forms for the mediation effects, are not easily generalizable to an outcome that is discrete or multivariate. In addition, model (2) does not permit a large number of mediators, e.g., more mediators than samples, unless some regularization is imposed.

### Inverse regression model

The limitations of the *forward* outcome model motivated us to adopt the *inverse* regression model that exchanges the positions of the outcome and mediators. Inverse regression is a commonly used approach, which, for example, has been widely used in genetics studies of multiple phenotypes [21–23]. It has a key advantage to allow different types of outcomes including multivariate outcomes. Also, it tends to offer other flexibilities, for example, allowing cell-type-specific mediation effects in testing the mediation of DNA methylation CpG sites [24]. Here we assume that a mediating taxon acts through its relative abundance, so *M*_*j*_ denotes the relative abundance of taxon *j*, although our methodology can easily accommodate presence-absence data (*M*_*j*_ takes 1 or 0 value indicating the existence of taxon *j* in a sample) without modification. We find that, by properly orthogonalizing the covariates against each other, we can obtain an inverse regression model that has the mediator model (1) “embedded” in. To this end, we define *T*_*r*_ to be the residual of *T* after orthogonalizing against *Z* and *O*_*r*_ the residual of *O* after orthogonalizing against (*Z, T*). We consider this inverse regression model:

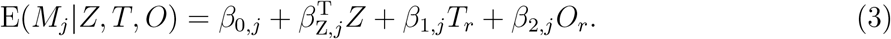

This model implies 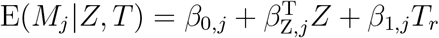, which is the mediator model (1) after orthogonalizing *T* against *Z*. Thus, *β*_1,*j*_ = *α*_1,*j*_. Although *β*_2,*j*_ ≠ *θ*_2,*j*_ due to inverse regression and marginal modeling of *M*_*j*_ in (3), we have that *β*_2,*j*_ = 0 and *θ*_2,*j*_ = 0 coincide. As a result, testing

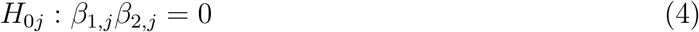

is equivalent to testing *α*_1,*j*_ *θ*_2,*j*_ = 0. We can test (4) by obtaining the least-squares estimates, 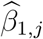 and 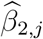, from (3), forming the test statistic 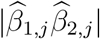, and using permutation to provide the null distribution of the test statistic. All of these can be achieved using the LDM with a slight modification on the test statistic.

Here we give a brief overview of the LDM. It was originally developed for testing bivariate associations between the microbiome and the covariates of interest. It is based on a linear model that regresses the microbiome data on the sequentially orthogonalized covariates that include first the confounding covariates that we wish to adjust for and then the covariates that we wish to test. The inference is based on permutation (i.e., permuting the orthogonalized covariates) to circumvent making parametric assumptions about the distribution of the microbiome data. The analyses of the raw relative abundance data and the arcsine-root transformed relative abundance data are performed, separately, and their results are combined to provide an omnibus test. The LDM allows an arbitrary number of taxa (including arbitrarily rare taxa) to be tested simultaneously with FDR control, supports continuous, discrete, or multi-variate covariates, accommodates clustered data [8], and offers analysis at the (transformed) relative abundance scale or the presence-absence scale [18, 19].

### Testing mediation effects at individual taxa

As mentioned after equation (4), it is most natural to consider the test statistic:

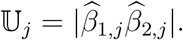

To provide a reference distribution for this statistic under the composite null of no mediation, we calculate the following statistic under the *b*th (*b* = 1,…, *B*) permutation:

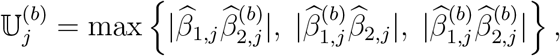

where 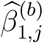 and 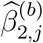 are obtained when *T*_*r*_ and *O*_*r*_ are permuted, separately, to break the *T* - *M*_*j*_ association given *Z* and the *M*_*j*_ -*O* association given (*Z, T*), respectively, and they are directly available in the LDM. The three product terms in 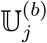 correspond to the test statistics under the type-I, type-II, and type-III null hypotheses. Thus, 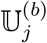 is inherently conservative in the sense that distribution is more spread out than the *true* distribution of 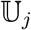 under a specific type of null (unknown). Finally, the permutation *p*-value for taxon *j* is calculated to be 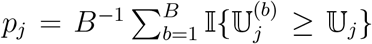, which is corrected for multiple testing by Sandve’s sequential stoping rule [25] as implemented in the LDM. We refer to this approach to testing individual taxa as LDM-med-product. However, it is unclear how to handle multivariate exposures or outcomes, in which case there are more than one element in *β*_1,*j*_ or *β*_2,*j*_.

A second way is to based the test statistic on the *p*-values *p*_1,*j*_ and *p*_2,*j*_ for testing *β*_1,*j*_ = 0 and *β*_2,*j*_ = 0, respectively, which naturally handle multivariate exposures or outcomes and are directly available from the LDM. We consider the test statistic:

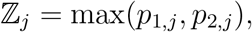

and assess the significance of ℤ_*j*_ by using the same permutation procedure as above and calculating the statistic:

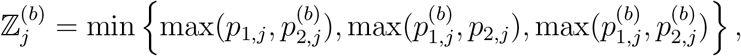

where 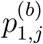 and 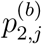 are based on the rank statistic of 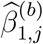 and 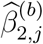, respectively, among all permutation replicates [26]. Note that max(*p*_1,*j*_, *p*_2,*j*_) can also be directly used as an analytical *p*-value for testing a single mediator [27], but here we choose permutation for inference because permutation is more robust and the permutation replicates are readily available from running the LDM. Similarly to 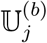, the statistic 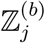 is conservative. Finally, the permutation *p*-value is calculated to be 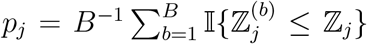 and corrected for multiple testing as in the LDM. We refer to this approach as LDM-med-maxP. In fact, this approach was found to be equivalent to LDM-med-product in simple settings, for example, when all variables are normally distributed [27]. However, both LDM-med-product and LDM-med-maxP tend to be inefficient for detecting significant mediators, which is a consequence of the conservative 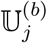 and 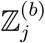 and the stringent correction over all *J* tests.

A third approach is to directly apply the MultiMed procedure [28] to *p*_1,*j*_ and *p*_2,*j*_, which was developed to improve efficiency of testing multiple mediators by restricting the testing to a subset of taxa that have relatively small *p*_1,*j*_ and *p*_2,*j*_. Here, we briefly describe this procedure; the theoretical properties that guarantee the FDR control can be found in the original papers [28, 29]. First, for a given significance level *α*, find the subset of taxa with relatively small *p*_1,*j*_ to be *ω*_*S*1_ = {*j*: *p*_1,*j*_ < *α*/2}, and denote the cardinality of the subset by 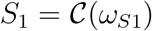. Similarly, find the subset with relatively small *p*_2,*j*_ to be *ω*_*S*2_ = {*j*: *p*_2,*j*_ < *α*/2} and denote 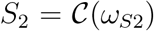. Then, the downstream testing of mediation is restricted to taxa at the intersection of the two subsets, which can greatly alleviate multiple testing correction. For taxon *j* ∈ *ω*_*S*1_ ∩ *ω*_*S*2_, define the subset-adjusted *p*-value:

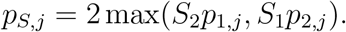

Taxon *j* is declared to be a mediator if the FDR-adjusted *p*-value

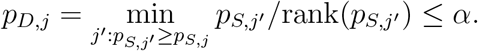

We call this approach LDM-med-subset. Although the subset-based approach has shown to be more efficient than the approach based on the product of coefficients (similar to our first approach) in the context of controlling the family-wise error rate [28], it is of interest to re-evaluate these approaches in the context of controlling the FDR.

### Testing the overall mediation effect in a community

If all taxa are under some types of null (not necessarily the same type), we declare a null community with no mediation effect. Given the *p*-values at individual taxa, it is straight-forward to construct a global test statistic by combining the individual *p*-values. Here we chose the *p*-value to be ℤ_*j*_ = max(*p*_1,*j*_, *p*_2,*j*_), and we adopt the Harmonic mean method [30] to aggregate ℤ_*j*_s, which is more robust to the dependence structure among taxa than Fisher’s method. The harmonic mean of ℤ_*j*_s is 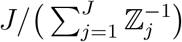, where a smaller value corresponds to a stronger evidence against the null hypothesis. To have a usual test statistic with a reverse directionality, we choose the statistic for the global test:

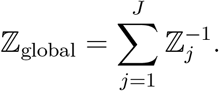

We assess the significance of ℤ_global_ via permutation, since permutation is more robust and the permutation replicates are readily available. The statistic based on *b*th permutation replicate is 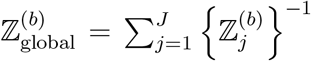. Finally, the permutation *p*-value for the global test is given by 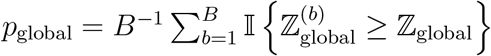. We call this test LDM-med-global, which is a natural extension of LDM-med-maxP but is also compatible with LDM-med-subset, which is also based on the association *p*-values *p*_1,*j*_ and *p*_2,*j*_.

## Results

### Simulation studies

Our simulations were based on data on 856 taxa of the upper-respiratory-tract (URT) microbiome [31] and the mediation model (1) and the forward outcome model (2) as generative models. Suppose that the exposure variable *T*_*i*_ is binary and that 50 samples were exposed (*T*_*i*_ = 1) and 50 unexposed (*T*_*i*_ = 0). We considered continuous outcomes as well as binary outcomes. We considered three mediation mechanisms, in which we assume the mediating taxa are the top five (1–5th) most abundant taxa, five (51–55th) less abundant taxa, and a selected mixture (4–5th and 51–52nd) of the two sets, and we refer to them as M-common, M-rare, and M-mixed. In all scenarios, we selected a set of taxa (6–10th) to be associated with the exposure but not the outcome, and another set of taxa (11–15th) to be associated with the outcome but not the exposure, corresponding to the type-I and type-II null taxa, respectively.

To generate the read count data, we first set the *baseline* (when *T*_*i*_ = 0) relative abundances of all taxa for all samples, denoted by 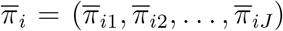, to the population means that were estimated from the real data. To induce the effects of the exposure *T*_*i*_ on a set of taxa (e.g., the mediating taxa or type-I null taxa), for those unexposed we kept 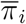 unchanged; for those exposed we decreased 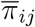 for some of the associated taxa by a percentage, which equals *β*_TM_ for the mediating taxa and *α*_TM_ (0 or 0.6) for the type-I null taxa, and we redistributed the decreased amount evenly over the remaining of the associated taxa; this way ensures that the relative abundances of non-associated taxa remain intact and the modified 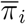 satisfies the compositional constraint (unit sum). Specifically, in M-common, the increasing set of the mediating taxa was taxa 1–2 and the decreasing set was taxa 3–5; in M-rare, the two sets were taxa 51–52 and 53–55; in M-mixed, the two sets were taxa 4 and 52 and taxa 5 and 51. Among the type-I null taxa, the two sets were taxa 6–7 and 8–10. The modified 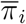 represents the *mean* relative abundances conditional on the exposure. Then, we introduced sample heterogeneity by drawing the sample-specific composition *π*_*i*_ = (*π*_*i*1_, *π*_*i*2_,…, *π*_*iJ*_) from the Dirichlet distribution 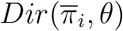 with mean 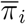 and overdispersion *θ* that was set to 0.02 (as estimated from the real data). Finally, we generated the read count data *M*_*i*_ = (*M*_*i*1_, *M*_*i*2_,…, *M*_*iJ*_) using the Multinomial distribution with mean *π*_*i*_ and the library sizes (sequencing depth) sampled from *N* (10000, (10000/3)^2^) and left truncated at 500.

To generate the outcome that is influenced by the mediating taxa, denoted by 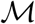, and the type-II null taxa, denoted by 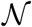, we partitioned each set of taxa into two subsets (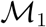 and 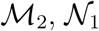 and 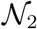) with approximately equal total relative abundance. In particular, we set 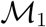 and 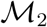 to be the increasing and decreasing sets, respectively, that were determined earlier relative to the exposure and have similar total relative abundance; we set 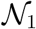 and 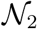 to be taxa 11–12 and 13–15, respectively. To simulate a continuous outcome, we used the model

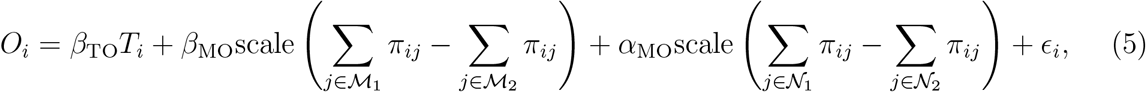

where scale(.) is a scaling function that standardizes a variable to have mean 0 and standard deviance 1, *β*_TO_ was fixed at 0.2, *α*_MO_ was fixed at 0 or 0.4, and the error term *ϵ*_*i*_ was drawn from *N* (0, 0.5^2^). It can be verified that the taxa that are neither mediators nor type-II null taxa were uncorrelated with the outcome after controlling for *T*_*i*_, owing to the counterbalancing effects of taxa in 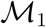 and 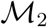 (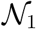 and 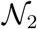) on the outcome. To simulate a binary outcome, we calculated the probability Pr(*O*_*i*_ = 1│*T*_*i*_, *π*_*i*_) = exp(*μ*_*i*_)/{1 + exp(*μ*_*i*_)} with *μ*_*i*_ being the same linear predictor as in (5), without the error term *ϵ*_*i*_.

To simulate a confounder, we note that a confounder has effects on the exposure, the microbiome, and the outcome (Figure 1(b)). Thus, we first simulated the binary confounder *Z*_*i*_ with 70% “success” rate among the exposed and 30% among the unexposed. Then, we used the same decreasing and increasing sets of the mediating taxa as determined earlier, now with the deduction percentage *γ*_ZM_ = 0.3, and the same operation as for the exposure to further modify 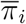 for those with *Z*_*i*_ = 1. Finally, we modified the linear predictor for the outcome to include the term 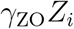 with *γ*_ZO_ = 0.7.

Prior to analysis, we filtered out taxa with fewer than 5 presence in the sample, which resulted in ~460 taxa remaining in analysis. For testing the overall mediation effect, we applied LDM-med-global and compared it to MedTest and MODIMA whenever the latter are applicable and valid. For MedTest, we adopted the omnibus test based on the Bray-Curtis and Jaccard distances, which focus on abundant and less abundant taxa, respectively, and thus form a complementary pair. For MODIMA, we chose Bray-Curtis, as it is the most commonly used distance in the literature and was also frequently used in the MODIMA paper. The type I error and power were assessed at the nominal level 0.05 based on 10000 and 1000 replicates of data, respectively. For testing mediation effects at individual taxa, we compared the three approaches: LDM-med-subset, LDM-med-maxP, and LDM-med-product. The sensitivity (proportion of the truly mediating taxa that were detected) and empirical FDR were assessed at the nominal FDR 10% based on 1000 replicates of data. Note that none of CMM, SparseMCMM, and Zhang’s method worked for our simulated data, as they either gave errors (due to the large number of taxa or extensive zero counts) or ran forever (> 10 hours).

### Simulation results

The type I error results of the global tests in M-common, M-rare, and M-mixed are summarized in Tables 1 and S1. We considered 12 scenarios under the global null hypothesis, each corresponding to a specific combination of the three types of null taxa. Clearly, MedTest and MODIMA easily lost control of the type I error in the presence of a mixture of the type-I and type-II null taxa. Note that the type-III null can be viewed as a special case of either the type-I or type-II null, so, for example, the first row in Table 1 corresponds to a global null where all taxa are under the type-II null. In all cases, LDM-med-global controlled the type I error; in fact, it was conservative as expected.

**Table 1:**
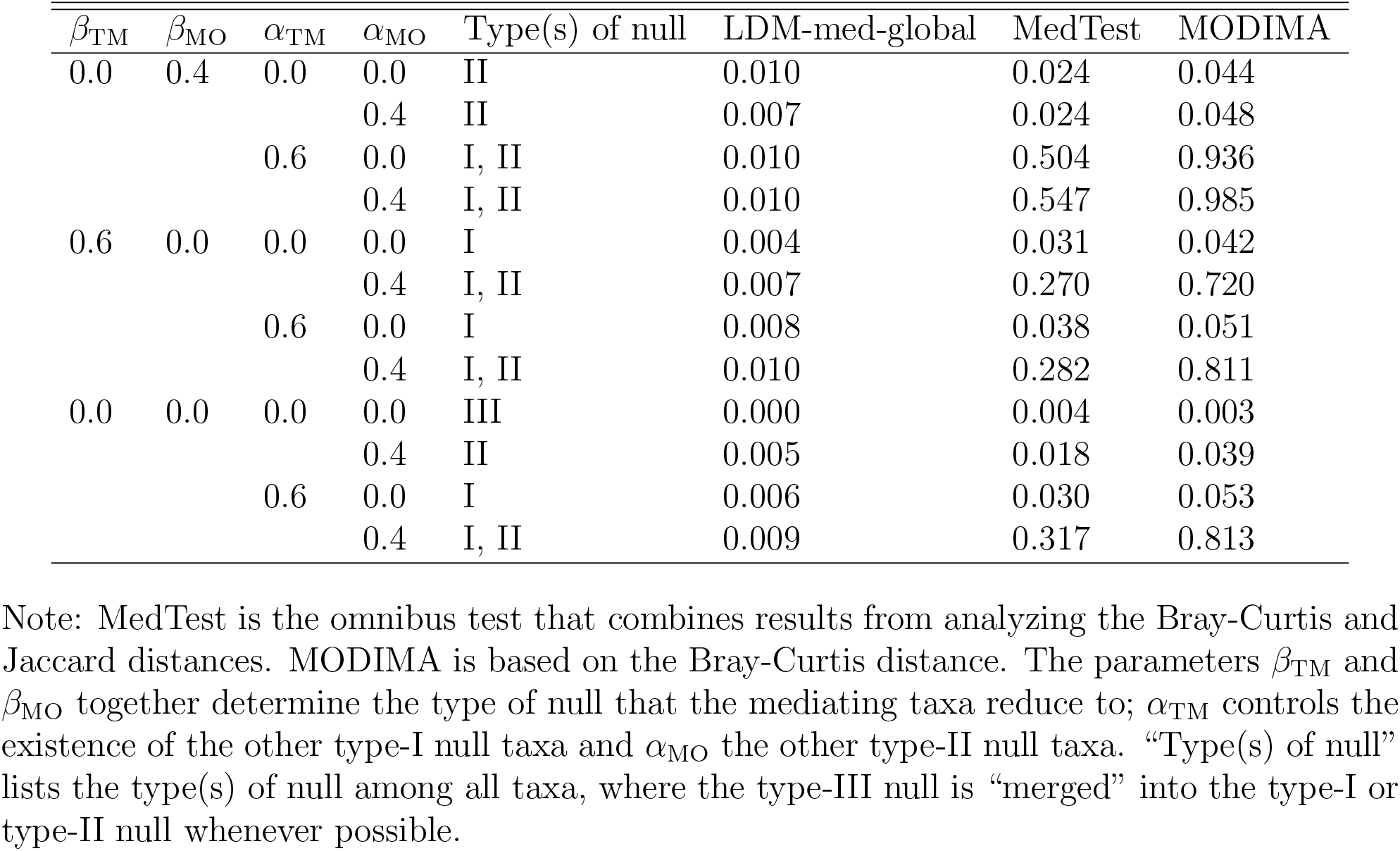
Type I error (at level 0.05) of the global tests in M-mixed with a continuous outcome and no confounder, in 12 scenarios under the global null

In the presence of a confounder (Table S2), LDM-med-global controlled the type I error even when the confounder was not adjusted for, due to its inherent conservativeness. This provides no clue to the extent of the confounding effect and the capability of LDM-med-global in adjusting for the confounding effect. Thus, we considered a variant of LDM-med-global, called LDM-med-global*, that used the information on the type of null for each taxon (only available in simulation) and set 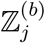 to the corresponding value among 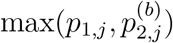 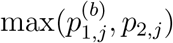 and 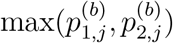. LDM-med-global*, yielded inflated type I error when the confounder was not accounted for and type I error close to 0.05 when it was accounted, demonstrating that LDM-med-global* (and hence LDM-med-global) was effective in adjusting for confounders.

For evaluating power, we started with the scenarios when there were neither type-I nor type-II null taxa (*α*_TM_ = *α*_MO_ = 0), under which MedTest and MODIMA were valid, and we also started with the simple case of a continuous outcome and no confounder. In M-common (Figure S1), MedTest and MODIMA were more powerful than LDM-med-global, whereas in M-rare (Figure S2), they were much less powerful than LDM-med-global, demonstrating that the two methods (with the most general choice of distance metrics) were effective in capturing variation in abundant taxa but not rare ones. In M-mixed (Figure 2), the power of LDM-med-global crossed with that of MedTest and MODIMA; LDM-med-global performed best when *β*_MO_ got relatively large. Then, we fixed the M-mixed scenario and varied other factors. When the confounder was introduced (Figure 3), the relative power of LDM-med-global and MedTest remained similar, in which case MODIMA was not included for comparison because it cannot adjust for additional covariates. When the type-I and type-II null taxa were introduced (Figure 4), they invalidated both MedTest and MODIMA but minimally affected the performance of LDM-med-global. When we switched to a binary outcome (Figure 5), LDM-med-global lost power to MedTest and MODIMA. We wished to know whether the power difference can be (partially) explained by the capability of LDM-med-global to allow different types of null at different taxa whereas MedTest and MODIMA cannot. To investigate this, we considered another variant of LDM-med-global, called LDM-med-global**, that assumed the same type of null (unknown) across all taxa as assumed in MedTest and MODIMA, and set 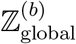 to 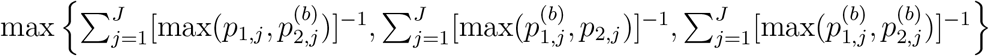. Indeed, LDM-med-global** gained substantially power over LDM-med-global and had comparable or even better power than MedTest. Note that LDM-med-global** is not restricted to binary outcomes.

**Figure 2:**
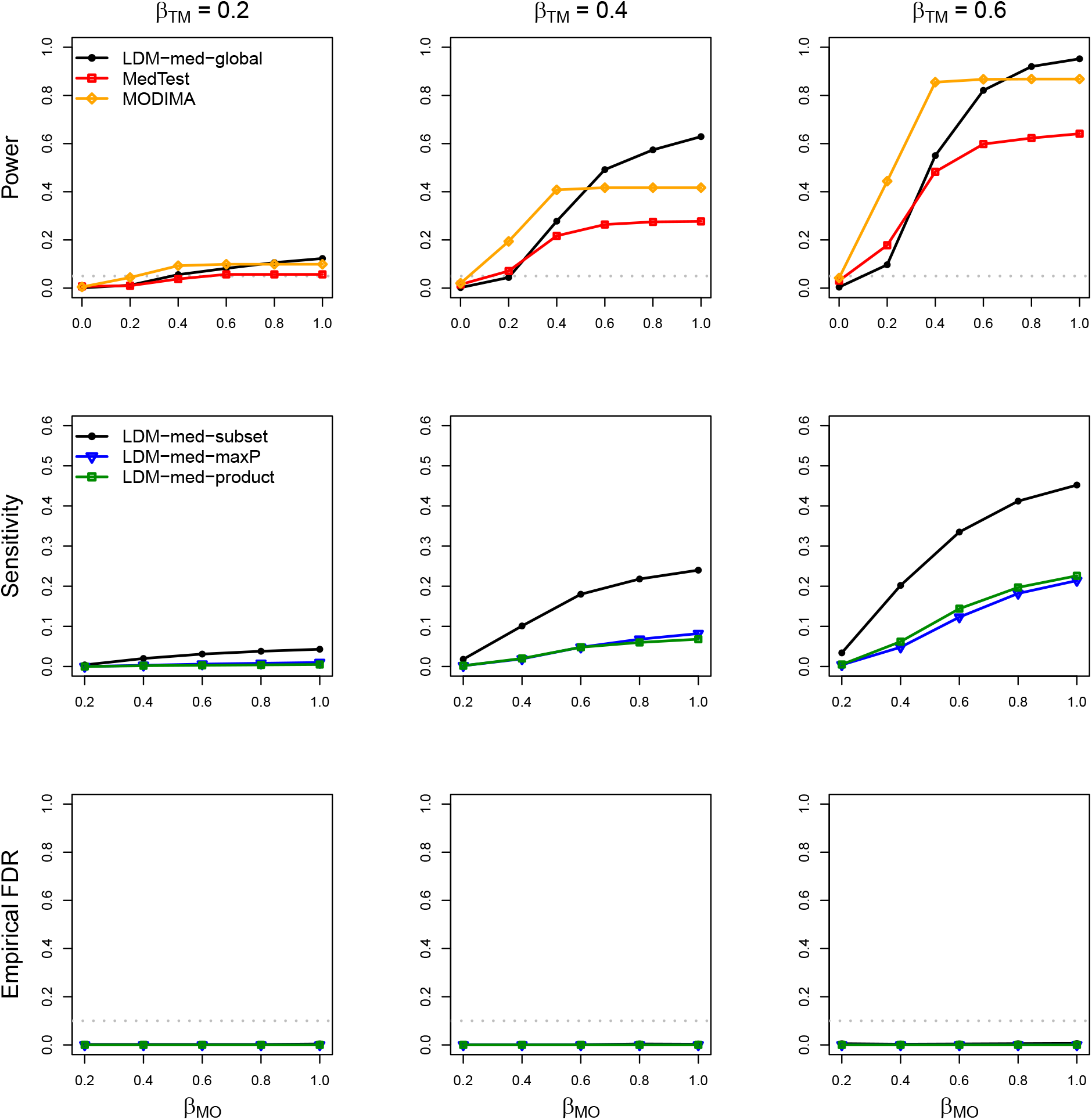
Simulation results in M-mixed with a continuous outcome and no confounder, in the absence of type-I and type-II null taxa (*α*_TM_ = 0 and *α*_MO_ = 0). The gray dotted lines represent the nominal levels 0.05 and 0.1 for power (or type I error when *β*_MO_ = 0.0) and empirical FDR, respectively.

**Figure 3:**
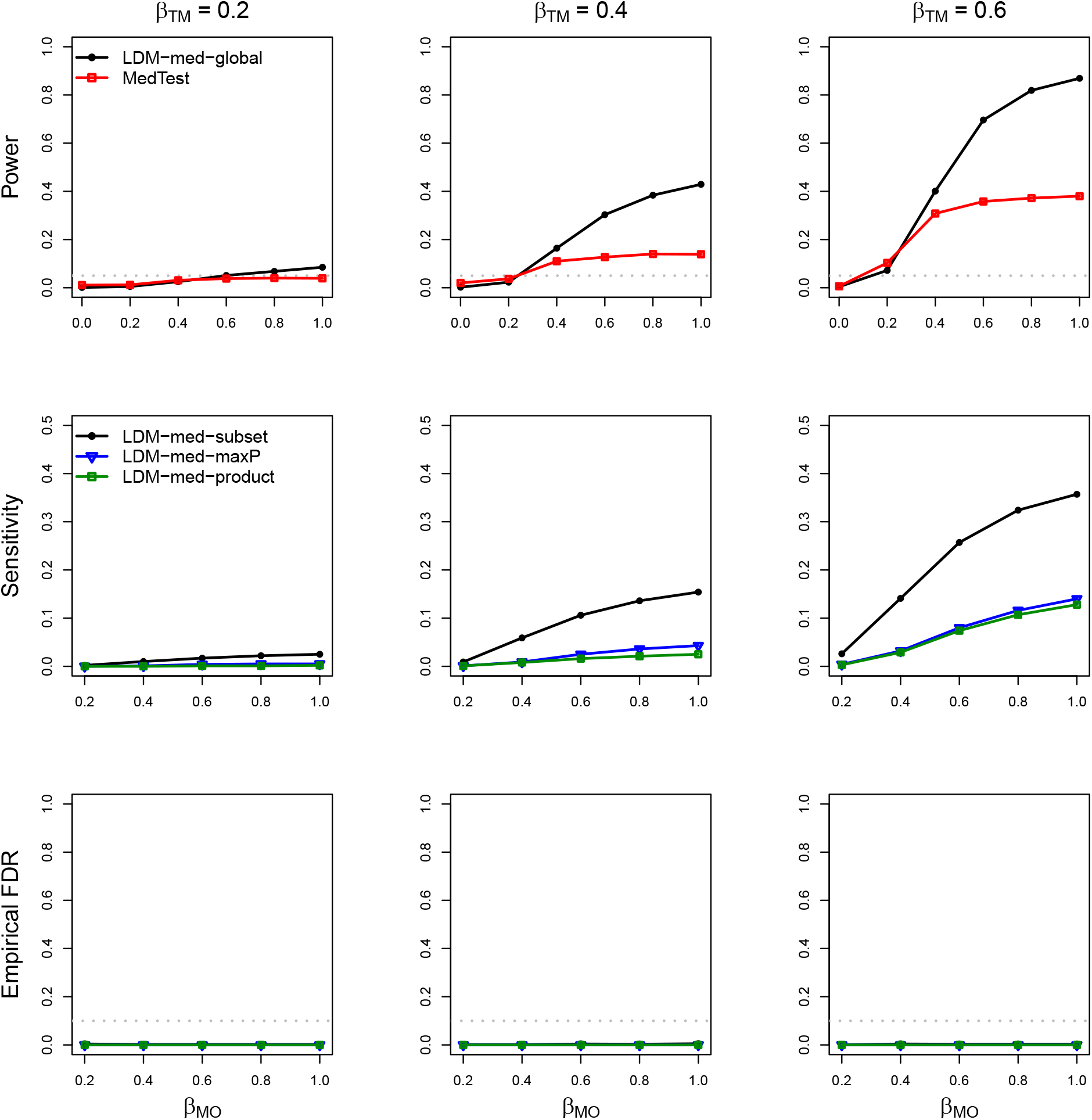
Simulation results in M-mixed with a continuous outcome and a confounder, in the absence of type-I and type-II null taxa. MODIMA was excluded because it does not allow adjustment of confounders.

**Figure 4:**
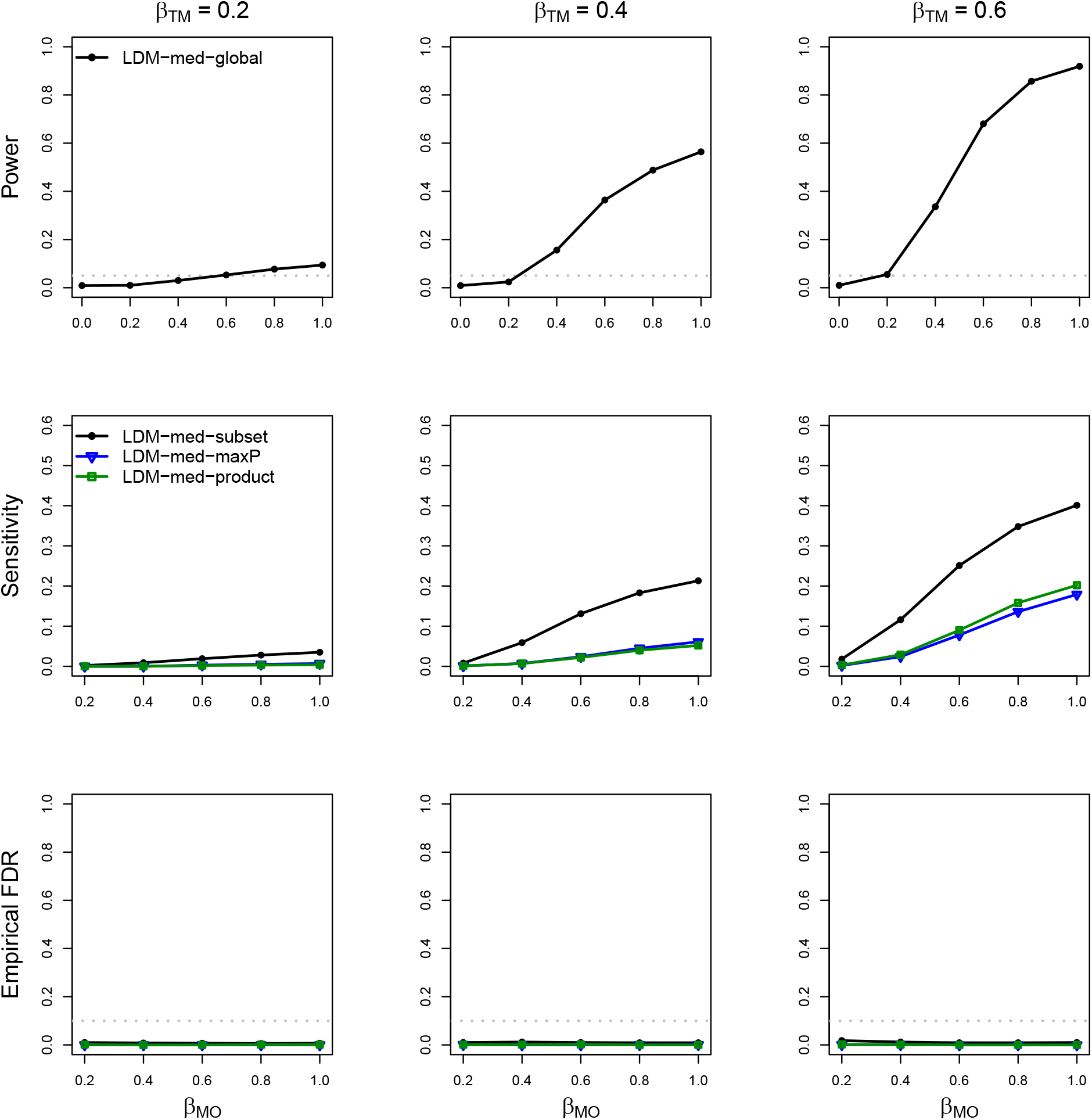
Simulation results in M-mixed with a continuous outcome and no confounder, in the presence of type-I and type-II null taxa (*α*_TM_ = 0.6 and *α*_MO_ = 0.4). MedTest and MODIMA were excluded because they did not control the type I error.

For testing mediation at individual taxa, all three LDM-based approaches controlled the FDR in all scenarios (Figures 2–5, S1–S2). As expected, the subset approach had substantially improved sensitivity over the product and maxP approaches, while the latter two had similar performance. For this reason, we chose LDM-med-subset as the method for testing individual taxa. We refer to the whole methodology as LDM-med, which consists of LDM-med-global and LDM-med-subset.

**Figure 5:**
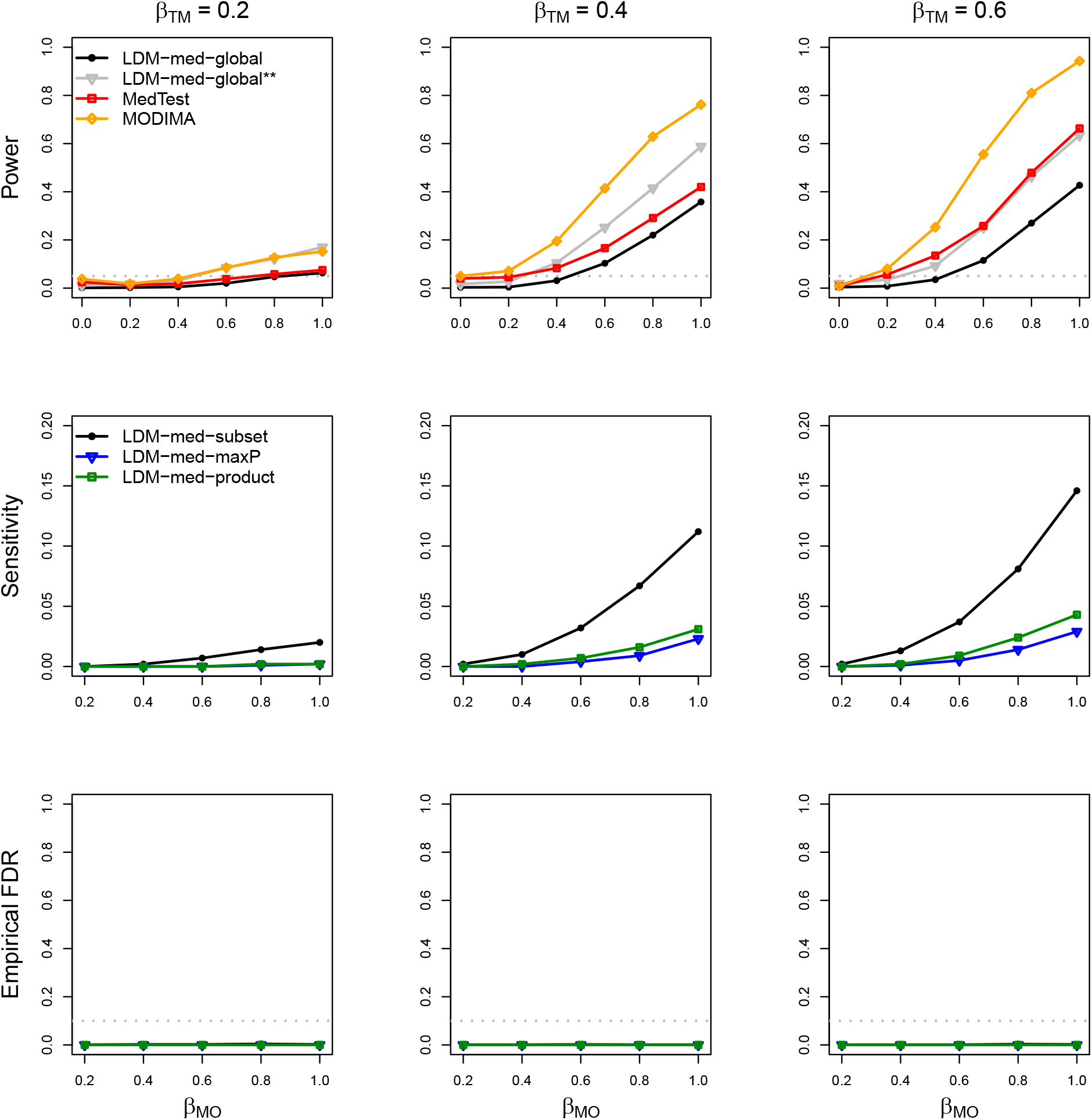
Simulation results in M-mixed with a binary outcome and no confounder, in the absence of type-I and type-II null taxa. LDM-med-global** is a variant of LDM-med-global that assumed the same type of null (unknown) for all taxa as was assumed in MedTest. The sample size was increased to 200 to obtain adequate power.

### Murine microbiome study

We analyzed the data generated from a murine microbiome study [32], which was conducted to explore whether the sub-therapeutic antibiotic treatment (STAT) would alter the gut microbiome composition and whether the shift of the gut microbiome would affect the body weight gain later in life. We focused on male mice for this analysis. The male mice were first randomized into the STAT and control groups, which was treated as a binary exposure variable. Then, their fecal samples were collected longitudinally at days 21 and 28. Bacterial DNAs were extracted from the fecal samples, sequenced for the 16S rRNA gene, and summarized into a taxa count table that initially contained 149 genera. Samples with less than 1800 reads, and genera with less than 10% presence or 0.01% mean relative abundance were filtered out, so the final taxa count table in our analysis included 41 genera and 36 mice (23 exposed to STAT and 13 unexposed) each having microbiome measurements at both time points. The mice body weight (in grams) prior to sacrifice was measured and used as a continuous outcome variable. There were no additional covariates to be adjusted for, as all potential confounders had been well-controlled in the randomized experiment.

It can be seen from Figure S3 that mice exposed to STAT were heavier than the control mice, with a small Wilcoxon *p*-value 0.011. We wish to know how much was this effect of STAT on body weight mediated through the gut microbiome. We tested the mediation effects at both the community level and the individual taxon level (at nominal FDR 10%) using LDM-med, MedTest, MODIMA, and SparseMCMM whenever they were applicable. Note that, although the outcome variable somewhat deviated from the normal distribution (Figure S3), all methods were robust because LDM-med treated the outcome as a covariate, and MedTest, MODIMA, and SparseMCMM based their inference on permutation.

We first restricted our mediation analysis to the cross-sectional microbiome data at day 28 only and summarized the main results in Table 2. LDM-med produced a global *p*-value 0.0351 and detected one significant mediator *[Ruminococcus]*. SparseMCMM yielded a more significant global *p*-value 0.004 and identified six mediators (with no FDR control though and no overlap with our detected mediator). Both MedTest (the omnibus test of Bray-Curtis and Jaccard distances) and MODIMA (based on the Bray-Curtis distance) produced non-significant global *p*-values 0.379 and 0.133, respectively, probably because *[Ruminococcus]* is rare with mean relative abundance 0.00089. To gain more insights into these results, we performed analysis of the bivariate association between the exposure and the relative abundance of each taxon using the Wilcoxon rank-sum test, and the bivariate association between the taxon and the outcome conditional on the exposure using the standard linear regression (treating the outcome as the dependent variable, and the exposure and taxon as the covariates), and corrected multiple testing in each association analysis by the Benjamini-Hochberg procedure at FDR 10%. As shown in Table S3, 22 taxa were found to be associated with the exposure, including *[Ruminococcus]* (unadjusted *p* = 0.012 and adjusted *p* = 0.019), and only *[Ruminococcus]* (unadjusted *p* = 0.0027 and adjusted *p* = 0.073) was found to be associated with the outcome. Thus, the mediator identified by LDM-med seems plausible.

**Table 2:**
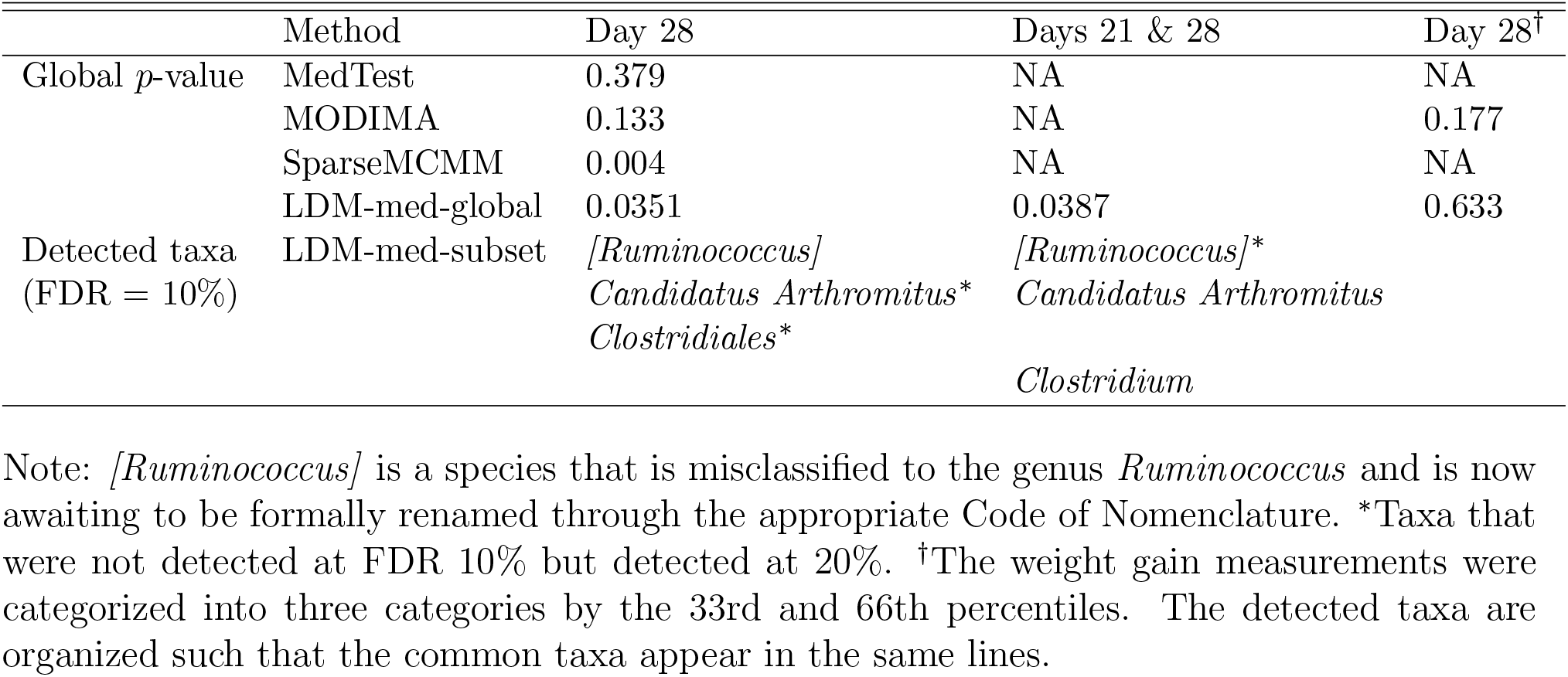
Mediation analysis of the murine microbiome study

We also performed mediation analysis of the longitudinal microbiome data at both days 21 and 28 and again summarized the main results in Table 2. Note that the outcome was observed only once per subject. While no other methods exist to analyze mediation of the microbiome data with correlations, LDM-med can easily handle such data because the LDM can handle clustered data (by setting `perm.within.type=“none”` and `perm.between.type=“free“`).

Here, a time variable (1/0) indicating day 28 was included as a covariate *Z*, as the microbiome composition was found to be significantly different between the two times (*p*-value 0.040 by the LDM for testing matched pairs). The results of mediation analysis by LDM-med were largely consistent with the previous results based on the data at day 28 only. The detected mediators based on the two datasets seemed non-overlapping at FDR 10% but substantially overlapped if the FDR was raised to 20%. We again performed analysis of bivariate associations between the exposure and each taxon using the LDM (adjusted for the time effect), and the taxon and the outcome conditional on the exposure using the standard linear regression (treating the outcome as the dependent variable, letting the covariates be the exposure variable, the relative abundances of the taxon at days 21 and 28, and testing the joint effect of the two relative abundance variables using the *F* test). The results were again largely consistent with the previous results on bivariate associations using the data at day 28 only (Table S3).

Finally, to illustrate the capability of LDM-med to handle categorical outcome variables, we converted the continuous outcome variable into three categories by the 33rd and 66th percentiles. For this type of outcome variables, only LDM-med and MODIMA were applicable, neither of which, however, identified any significant mediation effect.

## Discussion

We proposed a new approach to mediation analysis of the microbiome that is based on inverse regression and the LDM. Thus, our method LDM-med offers maximum robustness to the complex features in the taxa count data (i.e., high-dimensionality, sparsity, compositionality, and overdispersion), and provides extensive flexibility to accommodate various exposures and outcomes and study designs. Specifically, using the simulated and real data, we have demonstrated the capabilities of LDM-med to deal with null taxa under different types of null hypothesis of no mediation, binary and multivariate outcomes, clustered data with the exposure and outcome variables varying *between* the clusters, and confounding covariates. In addition, LDM-med can also handle clustered data with the exposure and/or outcome variables varying *within* the clusters [8], and perform analysis at the presence-absence scale using a rarefaction-without-resampling approach [18]. In summary, LDM-med can be highly useful in practice.

We have added LDM-med to our existing LDM R package. The computation of LDM-med is as efficient as the LDM. For example, using a single-thread MacBook Pro laptop (2.9 GHz Quad-Core Intel Core i7, 16GB memory), it took 46s to analyze one of our simulated datasets having 100 samples and ~460 taxa (after filtering). The murine dataset was at a smaller scale, consisting of 36 mice and 41 genera, so it took only 5s and 12s to analyze the data at day 28 only and the data at both day 21 and day 28, respectively.

LDM-med is based on marginal models for each mediator, and thus the identified mediators may not all be true biological mediators, which are called “probable mediators” but not “true mediators” [28]. This compromise was made in order to obtained controlled FDR for the selected mediators, which we deem as critical in the initial “scan” of high-dimensional features to generate “targets” to follow up in the downstream mechanistic study. This strategy has been very common in the analysis of high-dimensional omic data [15, 17, 28].

Our treatment of compositionality is different from that of CMM, SparseMCMM, and Zhang’s method, in that our method is based on the relative abundance data directly whereas the other methods are based on some type of log-ratio transformation of the relative abundance data. The two treatments correspond to two biological models for how microbial communities may change across conditions. The hypothesis in the other methods corresponds to the scenario that a small number of microbes have “bloomed” while the absolute counts of the others have not changed, which is commonly referred to as the “compositional” hypothesis. However, microbes interact with each other: not only do they compete for resources, but they also change their environment in ways that favor some microbes and suppress others. Because the microbiota are a community, it is not unreasonable to expect that many taxa changes when the condition changes. This hypothesis is referred to as the “community change” null hypothesis; the concepts “community state types” and “alpha diversity” exemplify this approach. When the “community change” hypothesis seems more reasonable, a method that applies directly to the relative abundance data such as the LDM and LDM-med is more appropriate.

## References

1. Bai J, Hu Y, Bruner D. Composition of gut microbiota and its association with body mass index and lifestyle factors in a cohort of 7–18 years old children from the American Gut Project. Pediatric Obesity. 2019;14(4):e12480.

2. Dunlop AL, Satten GA, Hu YJ, Knight AK, Hill CC, Wright ML, et al. Vaginal Microbiome Composition in Early Pregnancy and Risk of Spontaneous Preterm and Early Term Birth Among African American Women. Frontiers in Cellular and Infection Microbiology. 2021;11.

3. Pope JL, Tomkovich S, Yang Y, Jobin C. Microbiota as a mediator of cancer progression and therapy. Translational Research. 2017;179:139–154.

4. Dolan KT, Chang EB. Diet, gut microbes, and the pathogenesis of inflammatory bowel diseases. Molecular nutrition & food research. 2017;61(1):1600129.

5. Wang C, Hu J, Blaser MJ, Li H. Estimating and testing the microbial causal mediation effect with high-dimensional and compositional microbiome data. Bioinformatics. 2019;p. doi: 10.1093/bioinformatics/btz565.

6. Berg G, Rybakova D, Fischer D, Cernava T, Vergès MCC, Charles T, et al. Microbiome definition re-visited: old concepts and new challenges. Microbiome. 2020;8(1):1–22.

7. Hu YJ, Satten GA. Testing hypotheses about the microbiome using the linear decomposition model (LDM). Bioinformatics. 2020;p. bbtaa260, https://doi.org/10.1093/bioinformatics/btaa260.

8. Zhu Z, Satten GA, Mitchell C, Hu YJ. Constraining PERMANOVA and LDM to within-set comparisons by projection improves the efficiency of analyses of matched sets of microbiome data. Microbiome. 2021;9(1):1–19.

9. Zhang J, Wei Z, Chen J. A distance-based approach for testing the mediation effect of the human microbiome. Bioinformatics. 2018;34(11):1875–1883.

10. Hamidi B, Wallace K, Alekseyenko AV. MODIMA, a Method for Multivariate Omnibus Distance Mediation Analysis, Allows for Integration of Multivariate Exposure-Mediator-Response Relationships. Genes. 2019;10(7):524.

11. Sohn MB, Li H. Compositional mediation analysis for microbiome studies. The Annals of Applied Statistics. 2019;13(1):661–681.

12. Sohn MB, Lu J, Li H. A compositional mediation model for a binary outcome: Application to microbiome studies. Bioinformatics. 2021;.

13. Hu Y, Satten GA, Hu YJ. LOCOM: A logistic regression model for testing differential abundance in compositional microbiome data with false discovery rate control. bioRxiv. 2021;p. https://doi.org/10.1101/2021.10.03.462964.

14. Zhang H, Chen J, Li Z, Liu L. Testing for Mediation Effect with Application to Human Microbiome Data. Statistics in Biosciences. 2019;p. 1–16.

15. Asher JE, Lamb JA, Brocklebank D, Cazier JB, Maestrini E, Addis L, et al. A whole-genome scan and fine-mapping linkage study of auditory-visual synesthesia reveals evidence of linkage to chromosomes 2q24, 5q33, 6p12, and 12p12. The American Journal of Human Genetics. 2009;84(2):279–285.

16. Hu Y, Lin D. Analysis of untyped SNPs: maximum likelihood and imputation methods. Genetic epidemiology. 2010;34(8):803–815.

17. Hu YJ, Sun W, Tzeng JY, Perou CM. Proper use of allele-specific expression improves statistical power for cis-eQTL mapping with RNA-seq data. Journal of the American Statistical Association. 2015;110(511):962–974.

18. Hu YJ, Lane A, Satten GA. A rarefaction-based extension of the LDM for testing presence-absence associations in the microbiome. Bioinformatics. 2021;p. https://doi.org/10.1093/bioinformatics/btab012.

19. Hu YJ, Satten GA. A rarefaction-without-resampling extension of PERMANOVA for testing presence-absence associations in the microbiome. bioRxiv. 2021;p. https://doi.org/10.1101/2021.04.06.438671.

20. VanderWeele T, Vansteelandt S. Mediation analysis with multiple mediators. Epidemiologic methods. 2014;2(1):95–115. PMCID: PMC4287269.

21. O’Reilly PF, Hoggart CJ, Pomyen Y, Calboli FC, Elliott P, Jarvelin MR, et al. MultiPhen: joint model of multiple phenotypes can increase discovery in GWAS. PloS One. 2012;7(5).

22. Wu B, Pankow JS. Statistical methods for association tests of multiple continuous traits in genome-wide association studies. Annals of Human Genetics. 2015;79(4):282–293.

23. Majumdar A, Witte JS, Ghosh S. Semiparametric allelic tests for mapping multiple phenotypes: Binomial regression and Mahalanobis distance. Genetic Epidemiology. 2015;39(8):635–650.

24. Luo X, Schwartz J, Baccarelli A, Liu Z. Testing cell-type-specific mediation effects in genome-wide epigenetic studies. Briefings in Bioinformatics. 2021;22(3):bbaa131.

25. Sandve GK, Ferkingstad E, Nygård S. Sequential Monte Carlo multiple testing. Bioinformatics. 2011;27(23):3235–3241.

26. Westfall PH, Young SS. Resampling-based multiple testing: Examples and methods for p-value adjustment. John Wiley & Sons; 1993.

27. Boca SM, Sinha R, Cross AJ, Moore SC, Sampson JN. Testing multiple biological mediators simultaneously. Bioinformatics. 2014;30(2):214–220.

28. Sampson JN, Boca SM, Moore SC, Heller R. FWER and FDR control when testing multiple mediators. Bioinformatics. 2018;34(14):2418–2424.

29. Bogomolov M, Heller R. Assessing replicability of findings across two studies of multiple features. Biometrika. 2018;105(3):505–516.

30. Wilson DJ. The harmonic mean p-value for combining dependent tests. Proceedings of the National Academy of Sciences. 2019;116(4):1195–1200.

31. Charlson ES, Chen J, Custers-Allen R, Bittinger K, Li H, Sinha R, et al. Disordered microbial communities in the upper respiratory tract of cigarette smokers. PloS one. 2010;5(12):e15216. PMCID: PMC3004851.

32. Schulfer AF, Schluter J, Zhang Y, Brown Q, Pathmasiri W, McRitchie S, et al. The impact of early-life sub-therapeutic antibiotic treatment (STAT) on excessive weight is robust despite transfer of intestinal microbes. The ISME journal. 2019;13(5):1280–1292.

33. Benjamini Y, Hochberg Y. Controlling the false discovery rate: a practical and powerful approach to multiple testing. Journal of the royal statistical society Series B (Methodological). 1995;p. 289–300.

