## Supplemental Figures and Tables for "A New Approach to Testing Mediation of the Microbiome using the LDM"

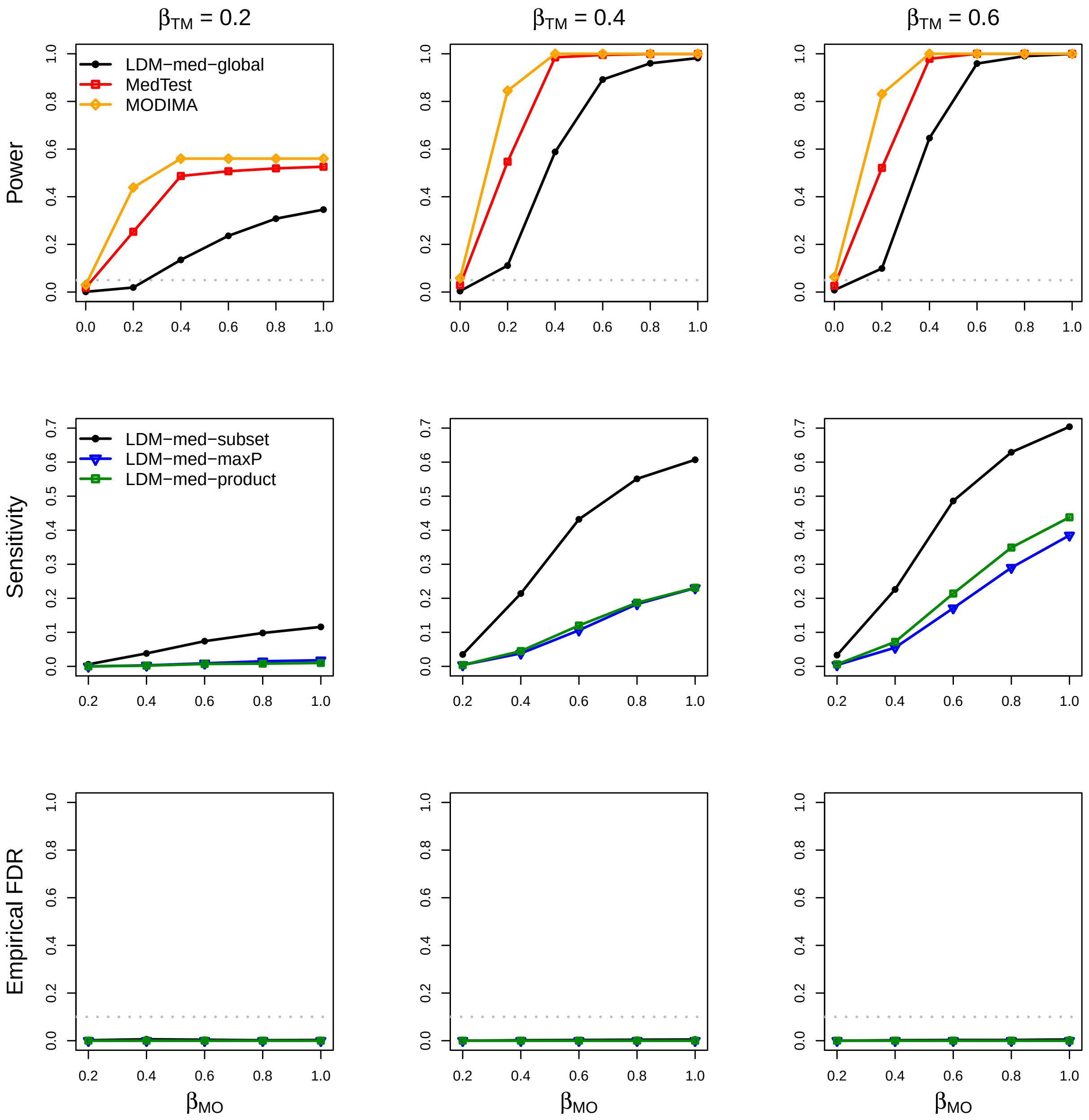


Figure S1: Simulation results in M-common with a continuous outcome and no confounder, in the absence of type-I and type-II null taxa.

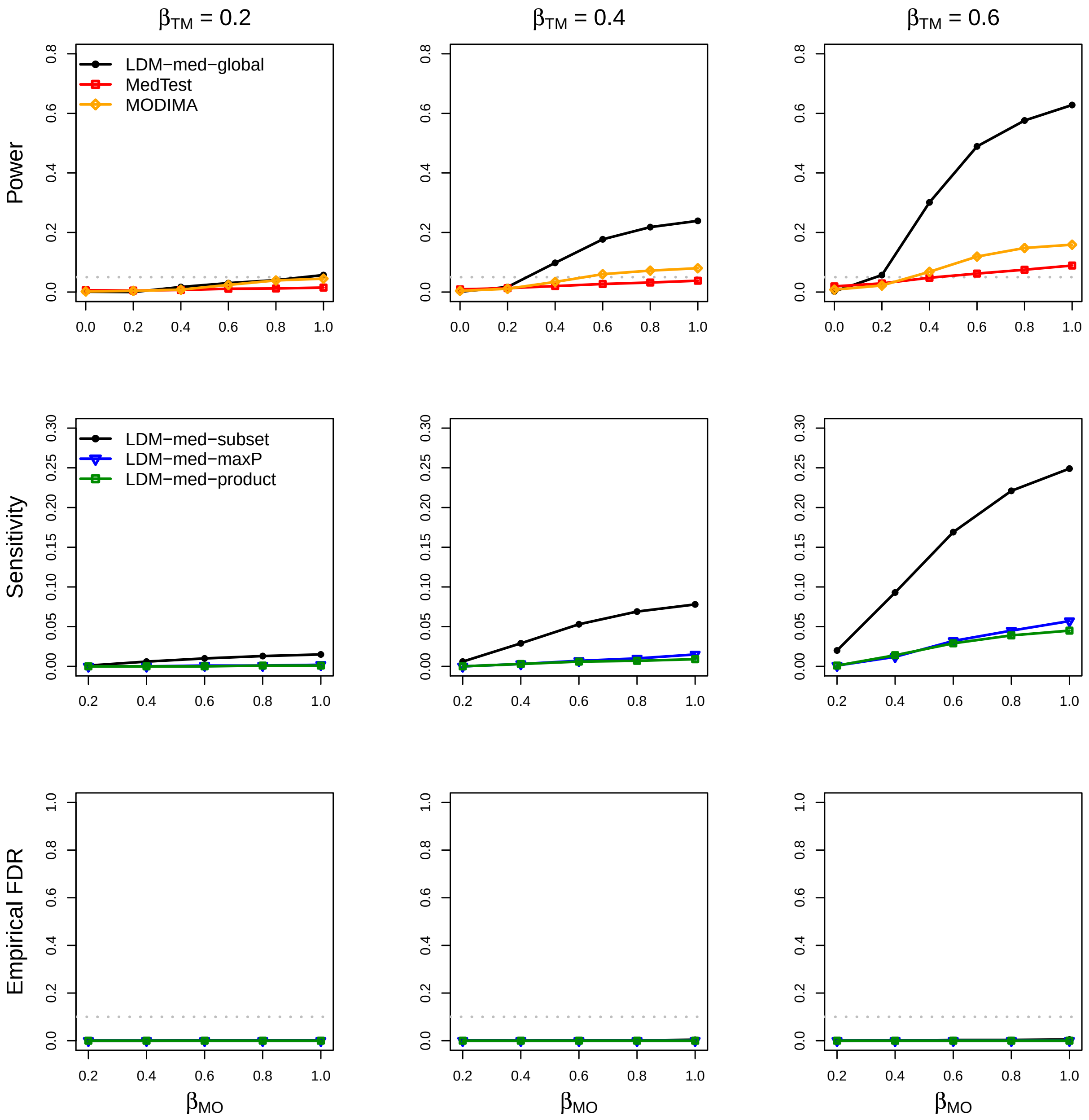


Figure S2: Simulation results in M-rare with a continuous outcome and no confounder, in the absence of type-I and type-II null taxa.

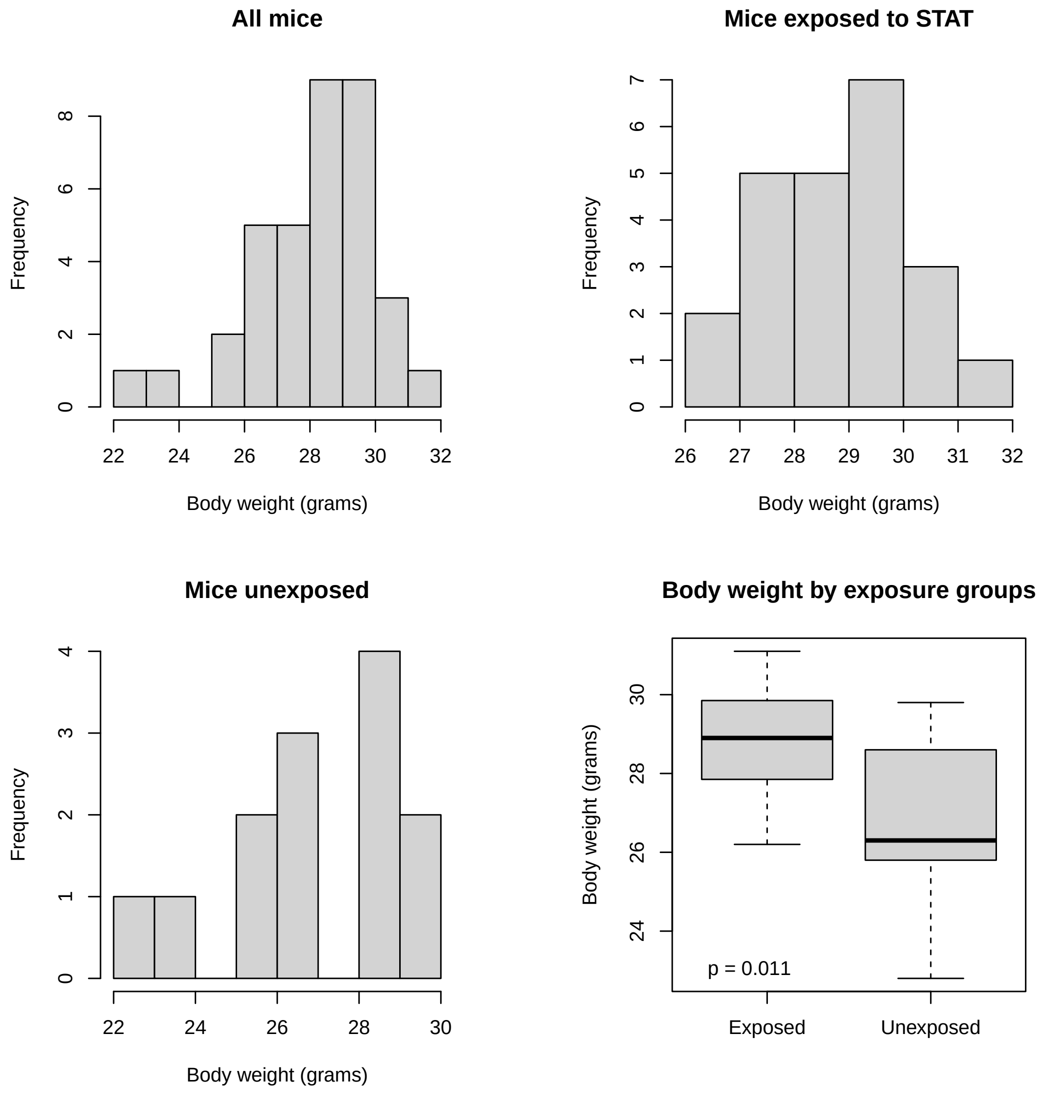


Figure S3: Distribution of the body weight outcome in the murine microbiome study.

Table S1: Type I error (at level 0.05) of the global tests in M-common and M-rare with a continuous outcome and no confounder, in 12 scenarios under the global null

| | $\beta_{\mathrm{TM}}$ | $\beta_{\mathrm{MO}}$ | $\alpha_{\mathrm{TM}}$ | $\alpha_{\mathrm{MO}}$ | Type(s) of null | LDM-med-global | MedTest | MODIMA |
|---|---|---|---|---|---|---|---|---|
| M-common | 0.0 | 0.4 | 0.0 | 0.0 | II | 0.004 | 0.028 | 0.048 |
| | | | | 0.4 | II | 0.005 | 0.028 | 0.048 |
| | | | 0.6 | 0.0 | I, II | 0.010 | 0.577 | 0.995 |
| | | | | 0.4 | I, II | 0.009 | 0.610 | 0.997 |
| | 0.6 | 0.0 | 0.0 | 0.0 | I | 0.008 | 0.026 | 0.063 |
| | | | | 0.4 | I, II | 0.010 | 0.102 | 0.767 |
| | | | 0.6 | 0.0 | I | 0.012 | 0.030 | 0.059 |
| | | | | 0.4 | I, II | 0.014 | 0.092 | 0.750 |
| | 0.0 | 0.0 | 0.0 | 0.0 | III | 0.000 | 0.004 | 0.003 |
| | | | | 0.4 | II | 0.005 | 0.018 | 0.039 |
| | | | 0.6 | 0.0 | I | 0.006 | 0.030 | 0.053 |
| | | | | 0.4 | I, II | 0.009 | 0.317 | 0.813 |
| M-rare | 0.0 | 0.4 | 0.0 | 0.0 | II | 0.002 | 0.010 | 0.013 |
| | | | | 0.4 | II | 0.003 | 0.020 | 0.039 |
| | | | 0.6 | 0.0 | I, II | 0.009 | 0.085 | 0.233 |
| | | | | 0.4 | I, II | 0.009 | 0.297 | 0.827 |
| | 0.6 | 0.0 | 0.0 | 0.0 | I | 0.002 | 0.019 | 0.008 |
| | | | | 0.4 | I, II | 0.006 | 0.055 | 0.139 |
| | | | 0.6 | 0.0 | I | 0.008 | 0.044 | 0.051 |
| | | | | 0.4 | I, II | 0.011 | 0.332 | 0.807 |
| | 0.0 | 0.0 | 0.0 | 0.0 | III | 0.000 | 0.004 | 0.003 |
| | | | | 0.4 | II | 0.005 | 0.018 | 0.039 |
| | | | 0.6 | 0.0 | I | 0.006 | 0.030 | 0.053 |
| | | | | 0.4 | I, II | 0.009 | 0.317 | 0.813 |

Note: see the Note to Table 1.

Table S2: Type I error (at level 0.05) of the global tests in M-mixed with a confounder and a continuous outcome, under three null scenarios

| | $\beta_{\text{TM}}$ | $\beta_{\text{MO}}$ | LDM-med-global | LDM-med-global* | MedTest |
|---|---|---|---|---|---|
| Adjusting for the confounder | 0.0 | 0.4 | 0.007 | 0.073 | 0.026 |
| | 0.6 | 0.0 | 0.004 | 0.069 | 0.020 |
| | 0.0 | 0.0 | 0.001 | 0.042 | 0.005 |
| Not adjusting for the confounder | 0.0 | 0.4 | 0.023 | 0.119 | 0.034 |
| | 0.6 | 0.0 | 0.016 | 0.108 | 0.032 |
| | 0.0 | 0.0 | 0.001 | 0.024 | 0.006 |

Note: we set $\alpha_{\text{TM}} = 0.0$ and $\alpha_{\text{MO}} = 0.0$. LDM-med-global* is a variant of LDM-med-global that used the information on the type of null for each taxa (only available in simulations). The type I error rates 0.073 and 0.069 after adjusting for the confounder were slightly inflated, due to the small sample size 100, and was reduced to 0.067 and 0.055 when the sample size was increased to 200.

Table S3: Bivariate association analysis of the murine microbiome study

| | Day 28 | Days 21 & 28 |
|---|---|---|
| Exposure–microbiome | | |
| Detected taxa (FDR = 10%) | *Candidatus Arthromitus* | *Candidatus Arthromitus* |
| | *Turicibacter* | *Turicibacter* |
| | *Clostridium.1* | *Clostridium.1* |
| | *RF39* | *RF39* |
| | *Dehalobacterium* | *Dehalobacterium* |
| | *Clostridiales* | *Clostridiales* |
| | *Ruminococcus* | *Ruminococcus* |
| | *Clostridiaceae* | *Clostridiaceae* |
| | *rc4-4* | *rc4-4* |
| | *Oscillospira* | *Oscillospira* |
| | *Dorea* | *Dorea* |
| | *[Ruminococcus]* | *[Ruminococcus]* |
| | *Allobaculum* | *Allobaculum* |
| | *Enterococcus* | *Enterococcus*$^{\dagger}$ |
| | *Lactobacillus* | |
| | *[Mogibacteriaceae]* | *[Mogibacteriaceae]* |
| | *Rikenellaceae* | |
| | *Erysipelotrichaceae* | *Erysipelotrichaceae* |
| | *Anaeroplasma* | |
| | *Clostridium* | *Clostridium* |
| | *Adlercreutzia* | *Adlercreutzia* |
| | *Coprococcus* | |
| | *Akkermansia*$^{*}$ | *Akkermansia* |
| | *Ruminococcaceae*$^{*}$ | |
| | | *Anaerostipes* |
| Microbiome–outcome \| exposure | | |
| Detected taxa (FDR = 10%) | *[Ruminococcus]* | *[Ruminococcus]*$^{\dagger}$ |

Note: $^{*}$Taxa that had $q$ values between 0.1 and 0.11; $^{\dagger}$Taxa that had $q$ values between 0.11 and 0.15. These taxa were considered as "nearly significant".
